# Pupil size asymmetries are modulated by an interaction between attentional load and task experience

**DOI:** 10.1101/137893

**Authors:** Basil Wahn, Daniel P. Ferris, W. David Hairston, Peter König

**Affiliations:** Institute of Cognitive Science, University of Osnabrück, Osnabrück, Germany; Human Neuromechanics Laboratory, School of Kinesiology, University of Michigan - Ann Arbor, MI, USA; Human Research and Engineering Directorate, U.S. Army Research Laboratory, Aberdeen, MD, USA; Department of Neurophysiology and Pathophysiology, Center of Experimental Medicine, University Medical Center Hamburg-Eppendorf, Hamburg, Germany

## Abstract

In a recently published study [1], we investigated how human pupil sizes are modulated by task experience as well as attentional load in a visuospatial task. In particular, participants performed a multiple object tracking (MOT) task while pupil sizes were recorded using binocular eyetracking measurements. To vary the attentional load, participants performed the MOT task either tracking zero or up to five targets. To manipulate the task experience, participants performed the MOT task on three consecutive days. We found that pupil sizes systematically increased with attentional load and decreased with additional task experience. For all these analyses, we averaged across the pupil sizes for the left and right eye. However, findings of a recent study [2] have suggested that also asymmetries in pupil sizes could be related to attentional processing. Given these findings, we further analyzed our data to investigate to what extent pupil size asymmetries are modulated by attentional load and task experience. We found a significant interaction effect between these two factors. That is, on the first day of the measurements, pupil size asymmetries were not modulated by attentional load while this was the case for the second and third day of the measurements. In particular, for the second and third day, pupil size asymmetries systematically increased with attentional load, indicating that attentional processing also modulates pupil size asymmetries. Given these results, we suggest that an increase in task experience (and associated reductions in arousal) uncover modulations in pupil size asymmetries related to attentional processing that are not observable for typical arousal levels. We suggest that these modulations could be a result of right-lateralized attentional processing in the brain that in turn influences structures involved in the control of pupil sizes such as the locus coeruleus. We can exclude a number of possible alternative explanations for this effect related to our experimental setup. Yet, given the novelty of this finding and the arguably speculative explanation of the underlying mechanisms, we suggest that future studies are needed to replicate the present effect and further investigate the underlying mechanisms.

## Introduction

For the past decade, researchers have investigated how modulations of pupil sizes are related to cognitive processes such as as decision-making [3–6], attention [7–21], emotions [22, 23], language [24], and memory [25–28]. Moreover, changes in pupil sizes have been repeatedly associated with changes in arousal [29–32]. In these studies, modulations of pupil size were either measured in only one eye or the pupil sizes were averaged across both eyes. However, to the best of our knowledge, only one study has also investigated whether pupil size asymmetries (i.e., differences in pupil sizes between the left and right eye) are related to attentional processing [2]. In this study [2], self-rated assessments of attention ability significantly correlated with pupil size asymmetries, suggesting that also pupil size asymmetries could be modulated by attentional processing. Such a link between pupil size asymmetries and attentional processing could be explained by the right-lateralization of attentional processing in the brain [33–35] and the relation of attentional processing to structures involved in the control of pupil sizes (i.e., the locus coeruleus [14, 32, 36–38]). That is, the right-lateralization of attentional processing could systematically affect structures related to pupil size control which in turn lead to a differential modulation of the left and right pupil sizes.

In a recent study [1], we investigated the relation between attentional demands, task experience, and pupil sizes (i.e., averaged across the left and right) in a visuospatial task (i.e., a multiple object tracking (MOT) task). In this study, to vary the attentional load, participants performed the MOT task either tracking zero or up to five targets. To manipulate the task experience, participants performed the MOT task on three consecutive days. We found that pupil sizes systematically increased with attentional load and decreased with additional task experience. However, to date, it has not been investigated whether changes in attentional load also differentially affect pupil size asymmetries. In order to address this question, we further analyzed the data from our previous study to investigate to what extent pupil size asymmetries are related to changes in attentional load. Given that we measured participants on three consecutive days, we also investigated the relation of pupil size asymmetries to task experience and the interaction between the factors attentional load and task experience.

## Results

For details on the methodology of the study, data preprocessing, behavioral performance in the MOT task, and results related to pupil sizes averaged across the left and right eye, we refer to our published manuscript [1].

For investigating pupil size changes that differentially affect the left and right eye, we performed the following normalization on the pre-processed data. We first subtracted the median pupil sizes of the right eye from pupil sizes of the left eye for each trial. Note, we took the median pupil size across three to nine seconds within the tracking period (i.e., while participants tracked the target objects on the computer screen) to avoid perceptual and executive confounds [1]. We then averaged these values for each participant, separately for each number of targets in the MOT task and day. We normalized the pupil size asymmetries for targets one to five by subtracting the pupil size asymmetries for the passive viewing condition and dividing the result by the pupil size in the passive viewing condition averaged across both eyes.

A descriptive overview of the normalized pupil size asymmetries can be seen in Fig 1. In this overview, no clear modulations of pupil size asymmetries by attentional load are visible on the first day. On the second and third day, however, pupil size asymmetries appear to be differentially affected by the attentional load conditions. This observation suggests an interaction effect between attentional load and task experience.

**Fig 1.**
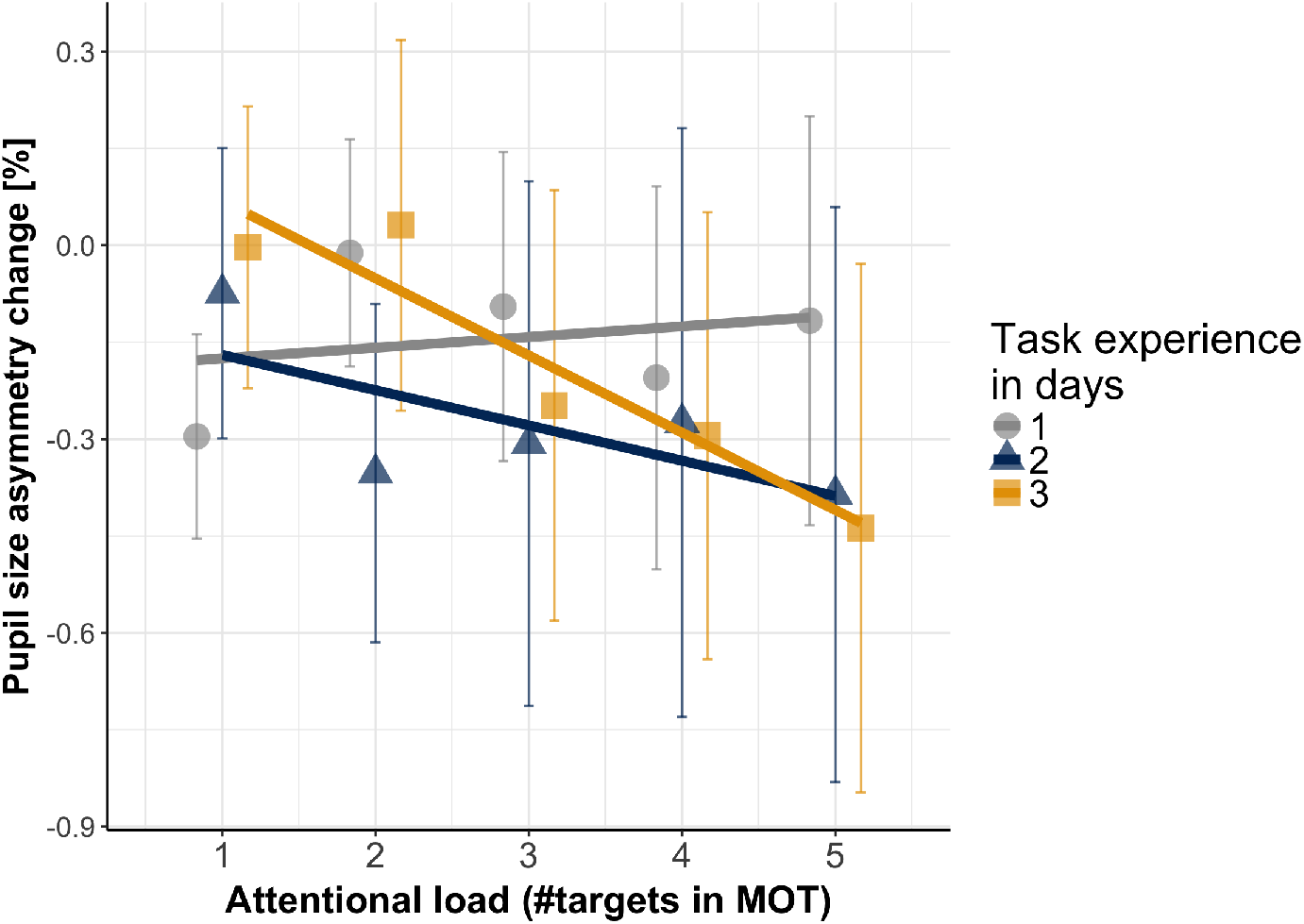
Pupil size asymmetries (left minus right) change relative to the passive viewing condition as a function of attentional load and task experience. Error bars are standard error of the mean. A linear regression fit is superimposed for each day.

For our inferential statistical tests, we used linear mixed models and model comparisons, starting with a baseline model in which we fitted intercepts as a random effect (i.e., intercepts were fitted for each participant individually) and used the pupil size asymmetries as the dependent variable. We then subsequently added attentional load (*χ*^2^(3)=34.06, *p* < .001, BIC = 841.1) and task experience (*χ*^2^(4)=85.27, *p* < .001, BIC = 778.7) as fixed and random effects to the model and both yielded significant results when performing model comparisons. Adding an interaction effect between attentional load and task experience as a fixed and random effect did yield a significant model comparison, suggesting that the effect of attentional load was modulated by the effect of task experience (*χ*^2^(1)=4.22, *p* = .040, BIC = 780.13). For the final model including all of these effects, we assessed the significance of the coefficients. We found a significant interaction effect (*B* = −0.07, *SE* = 0.03, *p* = .040) suggesting that with increasing task experience, the degree to which attentional load manipulations predict pupil size asymmetries increased. The coefficients for attentional load (*B* = 0.08, *SE* = 0.08, *p* = .326) and task experience (*B* = 0.25, *SE* = 0.19, *p* = .197) by themselves were not significant.

We followed up this analysis by calculating a separate Pearson correlation coefficient for each participant for each day, correlating the attentional load with the pupil size asymmetries. For this measure, a negative correlation would indicate that the pupil size asymmetries become larger with increasing attentional load. We tested these correlation coefficients against zero using a one sample t-test - for a descriptive overview, see Fig 2). Note, prior to entering the correlations in the t-test, we applied a Fisher z-transformation. These t-tests were not significant with regard to the first day (averaged *r* = -0.33, *t*(19) = 1.24, *p* = .230) and second day (averaged *r* = −0.26, *t*(19) = −1.60, p = .127)) but were significant for the third day (averaged *r* = −0.33, *t*(19) = −2.51, *p* = .021). These results suggest that at least for the third day, pupil size asymmetries are modulated by attentional load.

**Fig 2.**
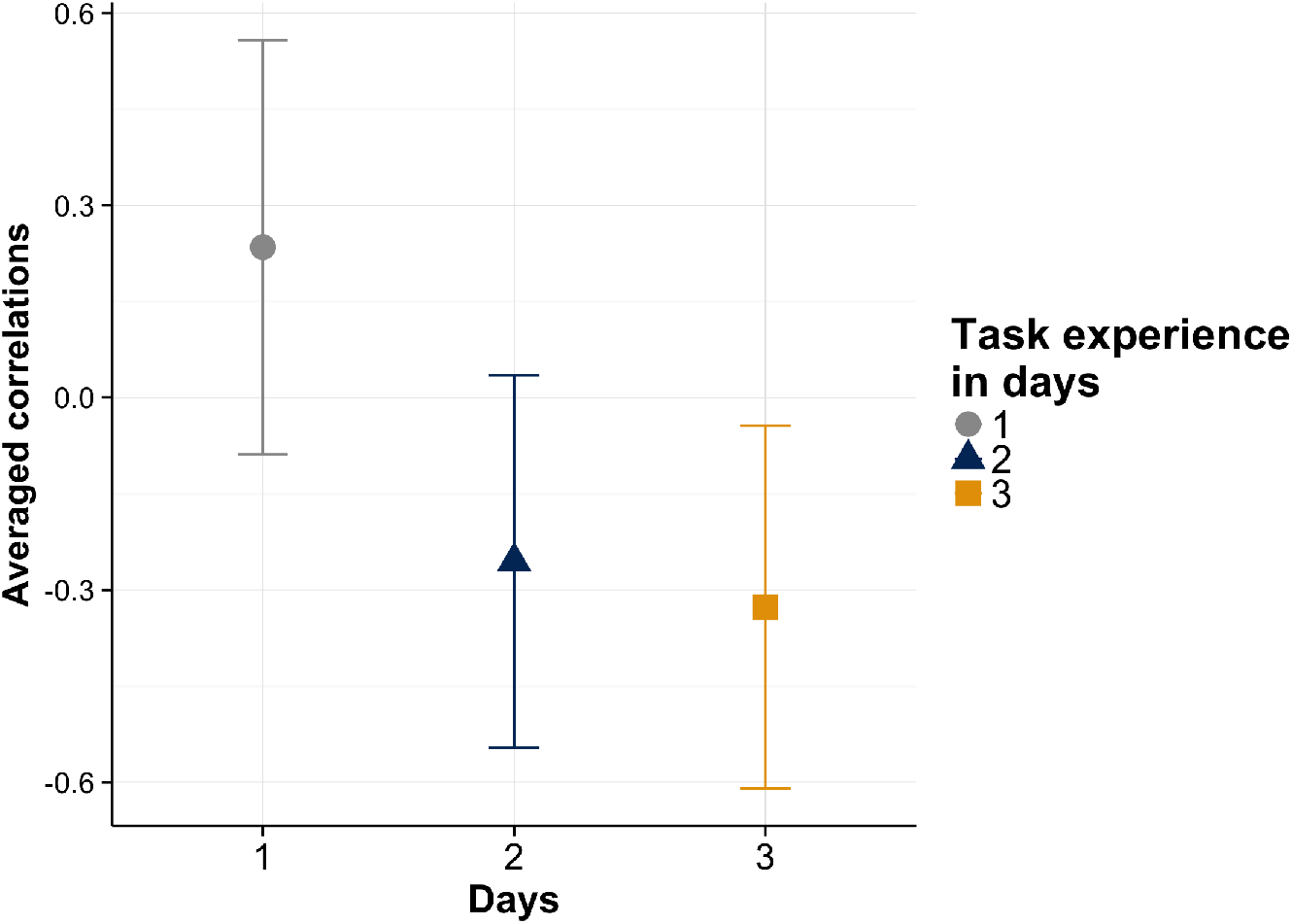
Averaged correlations (across participants) as a function of days. Error bars are confidence limits.

## Discussion

In sum, we found that pupil size asymmetries between the left and right eye are modulated by the attentional load in a visuospatial task. However, these modulations were small, and only present on the second and third day of the measurements. Moreover, these modulations were not as fine-grained as those observed for the pupil sizes averaged across both eyes [1].

Given that these modulations were only present on the second and third day of the measurements suggests that pupil size asymmetries are additionally influenced by task experience and possibly associated reductions in arousal. That is, larger effects modulating pupil sizes possibly due to arousal could mask smaller modulations related to pupil size asymmetries and attentional processing. Modulations of pupil size asymmetries related to attentional processing may only become visible after effects of arousal are considerably reduced.

Typically, asymmetries in pupil sizes are related to pathological conditions and are assumed to be consensual otherwise. In such pathological cases, the magnitude of the asymmetry is considerably larger than reported in the present study. That is, asymmetries are clearly visible when pupil sizes are visually inspected [39, 40]. Here, however, the reported data suggests that pupil size asymmetries also could be modulated by an interaction of attentional load and task experience. As pointed out in the introduction section, these modulations could be a result of right-lateralized attentional processing in the brain [33–35] that in turn influences structures involved in the control of pupil sizes such as the locus coeruleus [14, 32, 36–38].

As a point of note, one could suggest that modulations of pupil size asymmetries can alternatively be explained by factors related to the setup. One example is that participants could lean more towards the left during measurements or the left camera could be positioned more away from the eye than the right camera. We did not specifically control for these factors in our setup. However, if these factors would systematically alter results, we likely would already have observed pupil size asymmetries on the first day of the measurements. Yet, pupil size asymmetries were only present on the second and third day of the measurements. Also note, our measurement of pupil size asymmetries is a directional measure of the asymmetries. That is, we always subtracted the pupil size of the right eye from the pupil size of the left eye). We did not use an absolute measure of the asymmetries, i.e., we do not take the absolute value of the calculated differences. We suspect that any factors related to the setup that spuriously could produce pupil size asymmetries are subject to random processes (e.g., as noted above, an individual participant could lean more towards the right while another participant could lean more to the left). These random processes should likely affect absolute measures of the pupil size asymmetries. However, these processes should not systematically affect directional measures of pupil size asymmetries as it is unlikely that a large majority of participants were systematically measured differently on the first day compared to the second and third day of the measurements. Moreover, measurements for the different days for participants were often conduced on overlapping days across participants. For instance, participants who were measured on their second or last day of the measurements were measured on the same day as participants who were measured for the first time. That is, undesired changes in the setup that could have caused a modulation of pupil size asymmetries would have affected measurements in each experimental session. In summary, presently we do not have an indication that our results are based on an undesired confound in the setup.

More generally, future studies could test whether the pupil size asymmetries reported here become more pronounced when pupil sizes are measured for a more extended period of time than three consecutive days. Furthermore, it is an open question whether this effect generalizes to other types of tasks involving other forms of cognitive load. For instance, a memory task or an arithmetic task would be informative examples. Moreover, explanations on brain regions involved in pupil dilation control have primarily focused on describing general modulations of the pupil size [14, 32, 36–38]. Given these findings, future neurophysiological studies could also investigate the underlying processes that modulate pupil size asymmetries and may relate these to structural asymmetries in the brain [41, 42].

## Acknowledgments

We want to thank Lisa Steinmetz, Artur Czeszumski, and Ashima Keshava for their help with the data collection. We gratefully acknowledge the support by H2020 - H2020-FETPROACT-2014 641321 - socSMCs (for BW), ERC-2010-AdG #269716 - MULTISENSE (for PK), and the Cognition and Neuroergonomics Collaborative Technology Alliance ARL W91 1NF-10-2-0022 (for DPF and WDH).

